# Impaired expected value computations in schizophrenia are associated with a reduced ability to integrate reward probability and magnitude of recent outcomes

**DOI:** 10.1101/389551

**Authors:** Hernaus Dennis, Michael J. Frank, Elliot C. Brown, Jaime K. Brown, James M. Gold, James A. Waltz

## Abstract

**ABSTRACT:** *Background:* Motivational deficits in people with schizophrenia (PSZ) are associated with an inability to integrate the magnitude and probability of previous outcomes. The mechanisms that underlie probability-magnitude integration deficits, however, are poorly understood. We hypothesized that increased reliance on “value-less” stimulus-response associations, in lieu of expected value (EV)-based learning, could drive probability-magnitude integration deficits in PSZ with motivational deficits.

*Methods:* Healthy volunteers (*n*= 38) and PSZ (*n*=49) completed a reinforcement learning paradigm consisting of four stimulus pairs. Reward magnitude (3/2/1/0 points) and probability (90%/80%/20%/10%) together determined each stimulus’ EV. Following a learning phase, new and familiar stimulus pairings were presented. Participants were asked to select stimuli with the highest reward value.

*Results:* PSZ with high motivational deficits made increasingly less optimal choices as the difference in reward value (probability*magnitude) between two competing stimuli increased. Using a previously-validated computational hybrid model, PSZ relied less on EV (“Q-learning”) and more on stimulus-response learning (“actor-critic”), which correlated with SANS motivational deficit severity. PSZ specifically failed to represent reward magnitude, consistent with model demonstrations showing that response tendencies in the actor-critic were preferentially driven by reward probability.

*Conclusions:* Probability-magnitude deficits in PSZ with motivational deficits arise from underutilization of EV in favor of reliance on value-less stimulus-response associations. Consistent with previous work and confirmed by our computational hybrid framework, probability-magnitude integration deficits were driven specifically by a failure to represent reward magnitude. This work reconfirms the importance of decreased Q-learning/increased actor-critic-type learning as an explanatory framework for a range of EV deficits in PSZ.

## INTRODUCTION

Many people with schizophrenia (PSZ) suffer from a reduced tendency to engage in goal-directed behavior (1, 2), termed amotivation, or avolition. These deficits in motivation can contribute substantially to poor functional capacity and quality of life (3-5). In one explanatory framework, we have suggested that motivational deficits result from a specific impairment in the ability to precisely represent the value of an action or choice (expected value; EV) coupled with overreliance on “value-less” stimulus-response associations (6-8). Support for this computational account, however, comes from studies using reinforcement learning (RL) paradigms in which reward probability (the chance of obtaining a reward) solely determines EV. Importantly, other evidence indicates that PSZ (with motivational deficits) are *especially* impaired when EV estimation depends on the integration of reward magnitude (size) and probability.

One important line of evidence suggesting deficits in the representation of EV comes from the Iowa Gambling Task (IGT; (9)), in which participants select from four card decks with varying reward magnitude and probability. Performance deficits on this task, in PSZ, are driven by a reduced ability to integrate long-term outcome magnitude and probability (10-12). These deficits extend to contexts in which participants choose between earning a small certain reward or gamble for a larger reward (a “framing” task) (13). When reward magnitude and probability vary continuously, impaired EV computations in PSZ with motivational deficits are primarily driven by decreased sensitivity to reward magnitude (14). In contrast, PSZ can optimize task performance eventually if they learn about reward probability (15). Thus probability-magnitude integration deficits in PSZ (with motivational deficits) may be driven by impaired sensitivity to reward magnitude, and somewhat spared tracking of reward probability. To date, however, the computational mechanisms associated with an inability to integrate reward probability and magnitude into the EV of a choice has not been investigated. The goal of the current study was to use computational modeling to provide a mechanistic account of probability-magnitude integration deficits in PSZ.

There are several approaches to RL in computational models, which may in turn relate to different cognitive and neural mechanisms. While all RL models involve the reward prediction error (RPE) as a critical quantity that drives learning, the way in which the RPE is calculated and used to optimize behavior differs among classes. In the Q-learning framework (16), RPEs are used to update the EV of every action and choices are then executed directly based on the relative EV estimates. Computational models and neural data suggest that the dynamic representation of EV - specifically, the integration of reward magnitude and probability – crucially involves orbitofrontal cortex (OFC) (17-22). In the actor-critic framework (23), however, RPEs are not used to directly update choice EVs. Instead, the direct actor develops action propensities in a “value-less” space; choice preferences are only indirectly related to RPEs generated by the critic. The formation of such stimulus-response associations is thought to arise by gradual tuning of basal ganglia (BG) synaptic weights in response to dopaminergic RPEs (24, 25).

Although the differences between these two model classes may seem subtle, and both algorithms will generally produce adaptive learning, they can make categorically different predictions in particular scenarios. For example, since the actor only relies on choice propensities, given a novel choice between an option that had yielded gains and another one that had merely avoided losses, the actor will not exhibit a preference for the higher EV option if they had both given rise to positive RPEs in the critic. Using a hybrid computational model that allows for parametric mixing between Q- and actor-critic learning, we have previously shown that decreased reliance on the former and a relative increase in the latter can account for decreased gain-seeking behavior and overvaluation of contextual information in PSZ with motivational deficits (6, 7), which both result in a poor representation of EV. Thus, decreased reliance on EV and overutilization of stimulus-response associations serves as one computational framework of impaired goal-directed behavior in individuals with motivational deficits.

It is important, however, to test formal theories across a wide range of settings that are not limited to a single paradigm (7). Here, we hypothesized that the hybrid computational framework, via a reduction in Q-relative to actor-critic-type learning, can potentially account for abnormal probability-magnitude integration. First, because actor weights increase with each critic RPE, in stochastic environments they do not converge to any expected value, and as such they are overly influenced by frequency. Second, neural network models of OFC and basal ganglia (26) have suggested that dysfunction of the OFC can bias BG choices to be primarily driven by reward probability, whereas OFC integrity was needed to improve EV estimation by reward magnitudes. Similar probability patterns have been observed in algorithmic versions of this BG model, as in the Opponent Actor Learning (OpAL) architecture, a modified actor-critic based on physiological properties of the BG (25). We thus hypothesized that such a probability bias might be captured by our hybrid computational model in terms of an overreliance on actor-critic versus Q-learning.

In the study reported here, participants were presented with an RL task in which they learned to select stimulus options with the highest reward value. Pairs of stimuli differed in their reward probability and magnitude. In a subsequent transfer phase, old and novel pairs of stimuli were presented. Optimal performance in this phase of the task crucially depended on one’s ability to integrate reward probability and magnitude.

Through computational modeling and statistical analyses, we tested for the first time whether probability-magnitude integration deficits in PSZ (with motivational deficits) could be explained by underutilization of EV and/or overreliance on stimulus-response associations. In line with previous work (6, 7, 27), we predicted that deficits in the representation of EV should become more apparent as choices become easier. That is, we predicted that performance differences between PSZ and controls would be largest when the difference in EV between two competing stimuli was greatest (note that greater deficits with *increasing* difficulty would point to a more general learning impairment). Secondly, we expected that probability-magnitude integration deficits in PSZ, as well as reliance on reward probability versus magnitude in the entire sample, would correlate with the degree to which individuals relied on Q-versus actor-critic type learning. Finally, we expected probability-magnitude deficits and computational evidence thereof to correlate with the severity of motivational deficits.

## METHODS AND MATERIALS

### Participants

Forty-nine participants with a diagnosis of schizophrenia or schizoaffective disorder (PSZ) and thirty-eight healthy volunteers **(HV)** matched on age, gender, and ethnicity (Table 1) completed a probabilistic reinforcement learning task as part of a study approved by the institutional review boards of the University of Maryland and the Maryland State Department of Health and Mental Hygiene. Inclusion criteria, cognitive and clinical assessment details are reported in the Supplemental Text (including cut-offs for less severe motivational deficit [LMD} and more severe motivational deficit [MMD] subgroups). Written informed consent was obtained from all participants prior to the experiment.

**Table 1.**
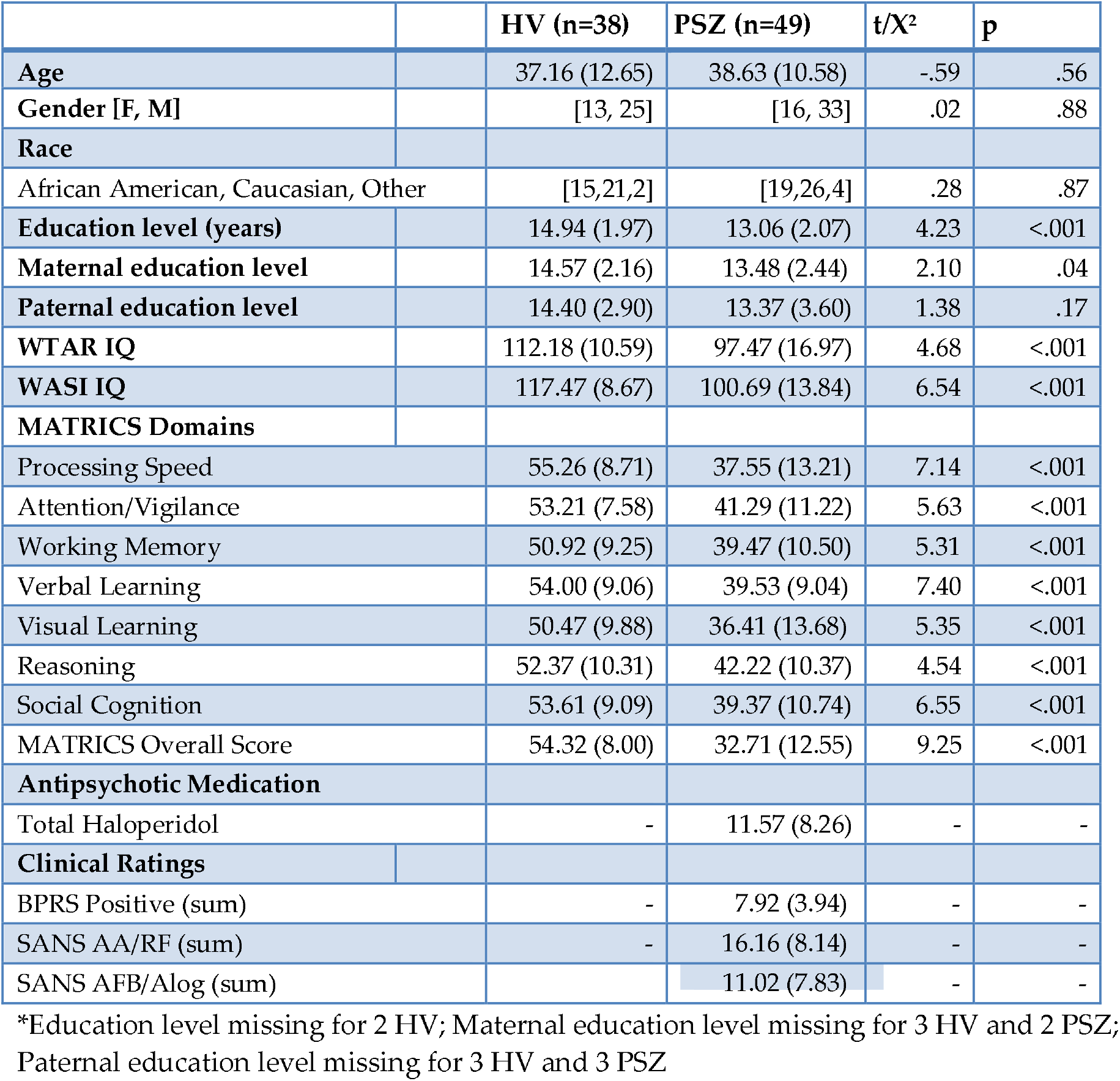
Sample Demographics.

### Probability-Magnitude Integration Task

Participants completed an RL task consisting of a Learning (160 trials) and a Test/Transfer (64 trials) Phase (Figure 1). During each acquisition phase trial, two stimuli were presented, on either side of a fixation cross. Participants were prompted to select one stimulus by pressing either the left or right trigger on a gamepad using their left or right index finger. Each choice was followed immediately by feedback, in the form of a number of points (+3, +2, +1, or +0). The eight stimuli differed in the probability and magnitude of the expected reward. For one pair, one stimulus resulted in a 3-point win on 90% of all trials, and a 0-point win on 10% of trials. The other option resulted in a 3-point win on 10% of all trials, and a 0-point win on 90% of trials (90-10/3 pair). The other three pairs were a “90-10/1” (1-point win 90% of the time for the optimal stimulus 1-point win 10% of the time for the non-optimal stimulus), “80-20/2” (2-point win 80% of the time for the optimal stimulus, 2-point win 20% of the time for the non-optimal stimulus) and “80-20/1” (1-point win 80% of the time for the optimal stimulus, 1-point win 20% of the time for the non-optimal stimulus) pair. All pairs were presented 40 times in pseudo-randomized order. Presentation order (optimal stimulus on left/right side of screen) was randomized across trials, and stimulus-value pairings were fully randomized across participants.

**Figure 1.**
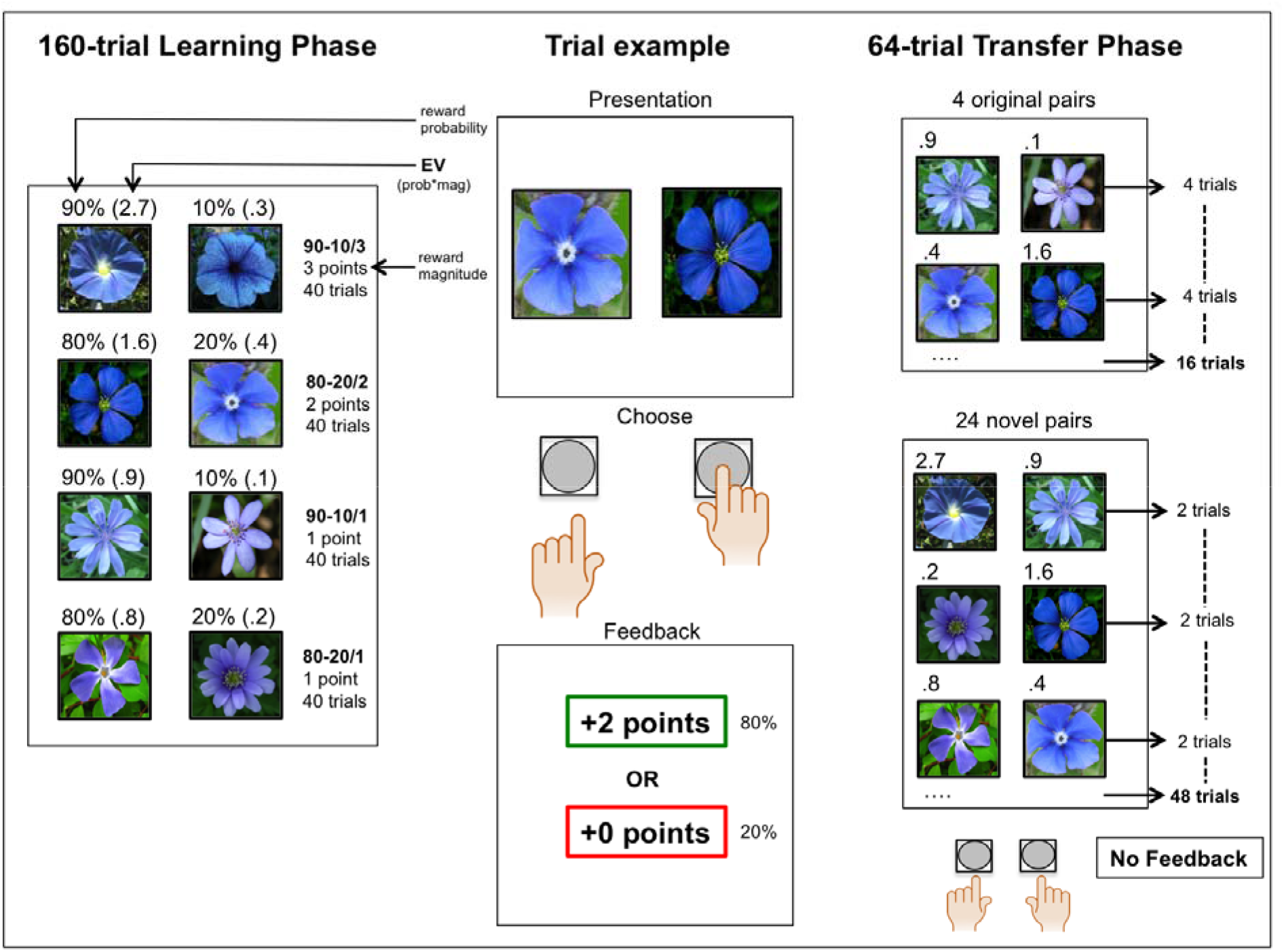
Graphical Overview Of The Probability-Magnitude Integration Task

The purpose of the Test/Transfer Phase was to assess participants’ ability to combine reward probability and magnitude into a representation of EV. Participants were presented with the four familiar Learning Phase pairs (“acquisition pairs”; 4 presentations per pair) and 24 novel pairs of stimuli (2 presentations per pair) and received the following instructions: “Please choose the picture that feels like it’s worth more points based on what you have learned during the previous block”. Crucially, for many of these trials, the optimal answer depended on the ability to combine the expected probability and magnitude of a stimulus (e.g. 80/2 vs. 90/1, or 10/3 vs. 20/2). No performance feedback was presented during the Test/Transfer Phase.

### Hybrid Computational Model

In order to represent the role of magnitude and probability in OFC- and BG-like computational frameworks, we adapted a previously-validated hybrid computational model that allows for mixed contributions from Q-learning and actor-critic frameworks (6, 7) (see “Model Selection and Comparison” in Supplemental Materials for details). In previous work, the frequency of RPEs was found to more efficiently update the formation of BG-like stimulus-response tendencies than the magnitude of the RPE (25, 26, 28, 29). This is related to the observation that frequency-driven RPEs, but not rare large magnitude-driven RPEs, drive choices in a neural network model of the BG (26). Accordingly, in “Model Selection and Comparison” (Supplemental Text) we demonstrate that the actor-critic component of our hybrid model is insensitive to the reward magnitude of outcomes. This is in line with BG-like algorithms, such as the actor-critic or OpAL architecture (24, 25), in which the formation of stimulus-response tendencies develops slowly with repeated sampling of stimuli. Therefore, large magnitude-driven RPEs are unlikely to rapidly counteract or enhance the effect of regular probability-driven RPEs. This is in contrast to models of OFC function, such as Q-learning, where RPEs directly update the EV, which in turn determines choice preferences. Here, magnitude-driven RPEs are used as effectively as probability-driven RPEs (26).

Using I) posterior predictive simulations (using the fitted parameters to simulate performance), II) demonstrations of learning on the basis of reward magnitude and probability, and III) quantifications of model fit, the “hybrid-probability” model was most likely to be the optimal model. Crucially, in the hybrid-probability model, Q-learning used both reward magnitude and probability, while the actor-critic only used the latter. The hybrid-probability model contained five free parameters: a critic (α_C_), actor (α_A_), and Q (α_Q_) learning rate, an inverse temperature parameter (*β*) that captured how deterministically participants sampled the optimal choice, and a mixing (m) parameter that weighted the contributions of Q- and actor-critic-type learning. Lower and upper bounds were set to 0 and 1 for all parameters. As *m* approximates 1, the contribution of Q-learning relative to actor-critic increases, while at *m*=.5 the contributions of both model classes are equal.

### Statistical Analyses

Effects of diagnostic group (PSZ vs. HV), trial block (4 blocks of 10 trials), and choice pair (90-10/3, 80-20/2, 90-10/1, 80-20/1) on acquisition pair performance were investigated using a repeated measures analysis of variance. Between-group differences in model parameters were ascertained using univariate analysis of variance and independent sample t-tests.

In accordance with previous work (7), novel stimulus pairs in the Test/Transfer Phase were ranked using the difference in EV between two competing stimuli (EV = probability*magnitude; Table S1 for details regarding stimulus combinations). A value difference tracking slope was computed for every participant using a logistic regression with value difference (value left – value right) as predictor and button press (left=1, right=0) as a nuisance regressor (thereby correcting for the tendency to select one side of the screen over the other).

Exploratory analyses into trials that were matched for reward magnitude while differing in reward probability (“probability discrimination trials”; 90/1-80/1), and trials that were matched for reward probability, while differing in reward magnitude (“magnitude discrimination trials”; 90/3-90/1 and 80/2-80/1), were conducted, as well as a group by trial-type (probability/magnitude discrimination) interaction. The latter analysis investigated whether performance advantages conferred by learning from reward magnitude versus learning from reward probability differed between the two diagnostic groups.

Correlation analyses were conducted using Spearman coefficients (due to non-normal distributions of many variables). Key findings, including group differences in Learning Phase accuracy, the value difference tracking slope, mixing parameter (HV vs. MMD) and correlations between the mixing parameter and Test/Transfer Phase performance in the entire sample, all survived Bonferroni correction for the number of model parameters (p=.01).

## RESULTS

### Demographics

Participant groups were matched on key demographic variables, including, age, gender, race, and paternal education, although PSZ scored lower on measures of IQ and all MATRICS subdomains (Table 1).

### Performance on Acquisition Pairs

In the Learning Phase, HV compared to PSZ overall made more optimal choices (F_(1,85)_=13.41, *p*<.001). There was no evidence for a group-by-trial block-by-pair (F_(7.22,613.92)_= .44, *p*=.88), group-by-pair (F_(2,68,228.01)_= .29, p=.81), or group-by-trial block (F_(2.33,197.73)_= .48, *p*=.77) interaction. HVs (trend-wise) outperformed PSZ on all stimulus pairs (90-10/3, t_(85)_=3.17, *p*=.002; 80-20/2; t_(85)_=2.90, *p*=.005; 90-10/1; t_(85)_=1.90, *p*=.06; 80-20/1; t_(85)_=2.62, *p*=.01; Figure 2A). Performance in block 4 (trials 31-40) was above chance in both participant groups for every pair (all *p*<.001).

**Figure 2.**
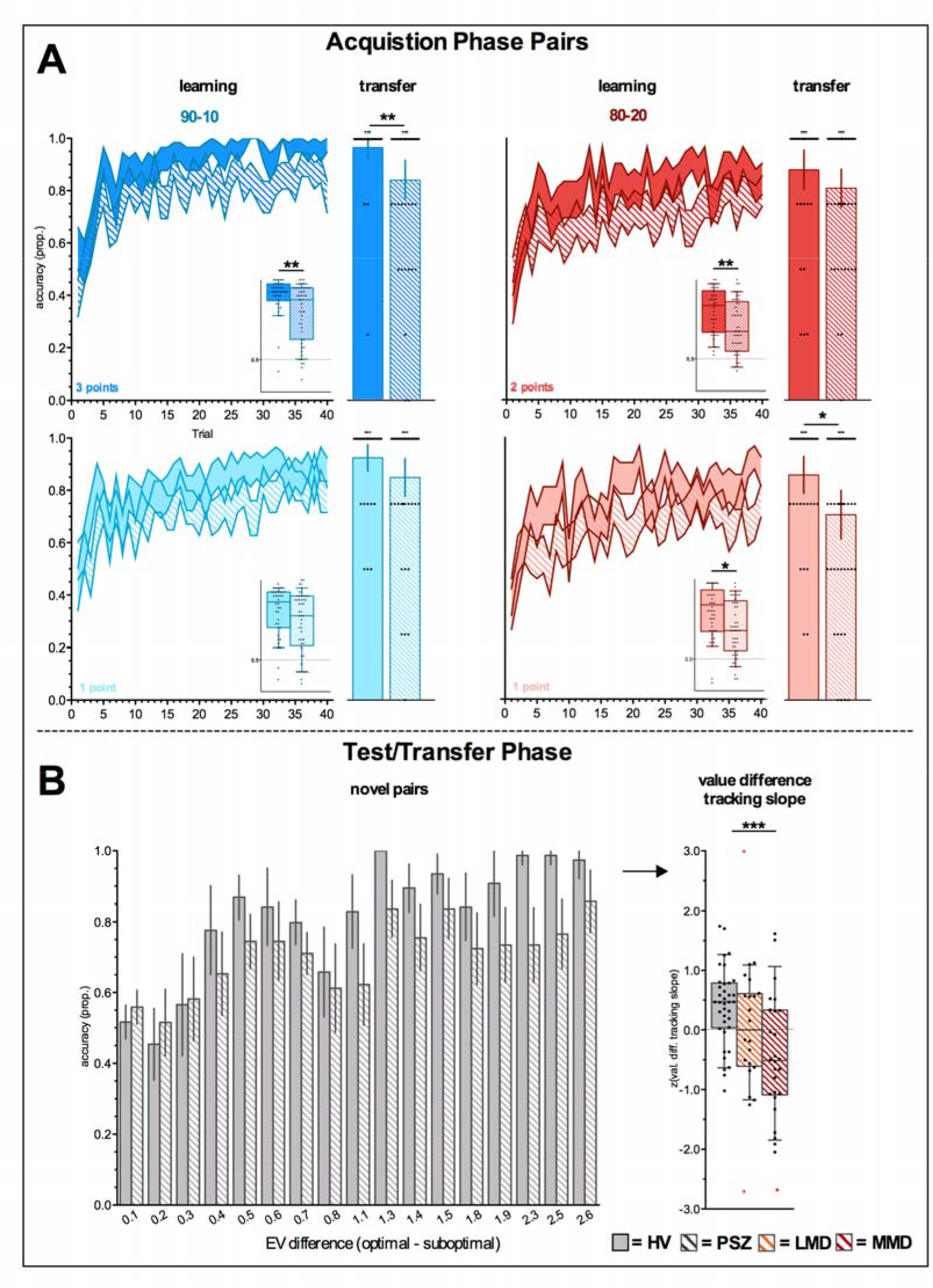
Performance on Acquisition Phase Pairs. **2A** Trial-by-trial (large figure) and average (small figure) Learning Phase accuracy (below “learning”), plotted next to average accuracy on acquisition phase pairs in the Test/Transfer Phase (below “transfer”). **2B** Test/Transfer phase performance on novel pairs ranked on EV (probability*magnitude) difference between two competing stimuli once presented for every value difference separately (“novel pairs”), once presented as the value difference tracking slope (for LMD and HMD separately; “value difference tracking slope”). 2 LNS and 1 HNS were removed from these analyses due to limited choice variability, which produced extreme performance slopes (marked in red). *p<.05, **p<.01, ***p<.001. Small asterisk above bars represents significance against chance.

In the Test/Transfer Phase, HV compared to PSZ also made more optimal choices (F_(1,85)_=10.26, *p*=.002). Specifically, HVs outperformed PSZ on 90-10/3 (t_(85)_=2.68, *p*=.009) and 80-10/1 (t_(85)_=2.51, *p*=.01), but not 80-20/2 (t_(85)_=1.34, *p*=.19) and 90-10/1 (t_(85)_=1.64, *p*=.10) trials (Figure 2A). Nevertheless, PSZ and HVs performed above chance on all original pairs (all *p*<.001), suggesting that both groups had developed a preference for the optimal stimulus.

Subgroup analyses in LMD and MMD revealed no evidence for an effect of motivational deficit severity on acquisition pair accuracy during either experiment phase (Supplemental Results; Figure S6).

### Performance on Novel Transfer Pairs

The value difference tracking slope was greater for HV than PSZ (t_(82)_=3.34, *p*=.001; 3 PSZ were removed due to limited choice variability; Figure S7 for performance on every Test/Transfer Phase pair for each diagnostic group). In line with predictions, these data indicate that PSZ improved less as the difference in EV between two competing options increased. Importantly, the group difference in the value difference tracking slope was driven by motivational deficit severity (HVs vs. LMD t_(56)_=1.78, *p*=.08; HVs vs. MMD t_(62)_=3.80, *p*<.001, LMD vs. MMD t_(44)_=.65, *p*=. 10; Figure 2B). These results suggest that MMD specifically were poorer at integrating reward probability and magnitude. BPRS positive symptom scores were not associated with the value difference tracking slope (Spearman’s rho=.15, *p*=.32)

Focusing on selective trials matched for probability and magnitude, we observed a group-by-trial type interaction (F_(1,85)_=4.35, *p*=.04; Figure 3). Post-hoc analyses revealed that HVs performed better on magnitude discrimination than probability discrimination trials (t_(37)_=3.59, *p*=.001), while PSZ performed similarly on magnitude and probability discrimination trials (t_(48)_=.73, *p*=.47) (Figure 3). The difference between performance on magnitude- and probability-discrimination trials - that is, the difference between the advantage conferred by higher reward magnitude versus higher reward probability-highly correlated with the value difference tracking slope, suggesting that participants that performed better on magnitude discrimination trials overall performed better in the Test/Transfer Phase (Spearman’s rho=.56, *p*<.001).

**Figure 3.**
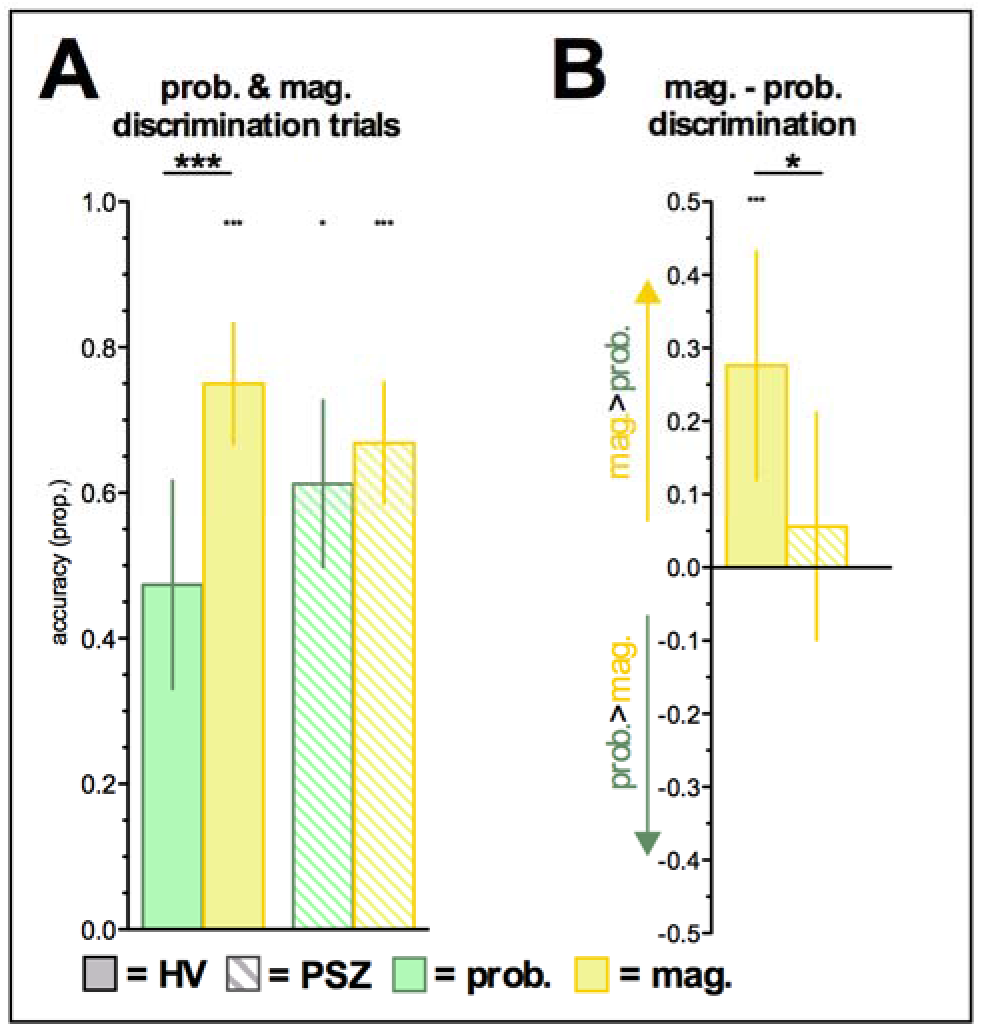
Performance on magnitude and probability discrimination trials during the Test/Transfer Phase for each diagnostic group. 3A Performance on probability (90/1-80/1) and magnitude (90/3-90/1 and 80/2-80/1) discrimination trials. 3B The difference in performance on magnitude and probability trials. *p<.05, ***p<.001. Small asterisk above bars represents significance against chance.

### Computational Modeling Analyses: Avolitional PSZ overutilize stimulus-response associations

Mean hybrid-probability model parameter estimates for each diagnostic group are reported separately in Table 2, with only the mixing parameter differing between HVs and PSZ (t_(85)_= 2.04, *p*=.04; Table S2 for individual model parameters), thereby replicating our previous results in a different paradigm setting (6, 7). Crucially, the group difference in the mixing parameter was driven by motivational deficit severity (HVs vs. LMD t_(58)_=.70, *p*=.48, Cohen’s *d*=. 19; HVs vs. MMD t_(63)_=2.69, *p*=.009, Cohen’s *d*=.67; LMD vs. MMD t_(47)_=1.55, *p*=. 13, Cohen’s *d*=.44; Figure 4A) and correlated with SANS total severity (Spearman’s rho=-.29, *p*=.04, df=48). The low mean mixing parameter (.40) suggests that MMD over-utilized actor-critic-type learning, while LMD (.57) and HVs (.64) relied more on Q-learning. We also observed a marginally significant difference in α_c_ (t_1,63_=1.99, *p*=.052), when we compared HV and MMD. This suggests that PSZ with motivational deficits updated their state value less in response to recent outcomes. Importantly, motivational deficit severity was not associated with model fit (SANS total score Spearman’s rho=-.11, *p*=.43; SANS AA score Spearman’s rho=-.13, *p*=.36). Moreover, BPRS positive symptom scores were not associated with the mixing parameter (Spearman’s rho=-.03, *p*=.82)

**Table 2.** Hybrid-probability model parameters.

|  | HV (n=38) | PSZ (n=49) | t | p |
| --- | --- | --- | --- | --- |
| Critic learning rate ($\alpha_c$) | .49 (.42) | .32 (.37) | 1.95 | .35 |
| Actor learning rate ($\alpha_a$) | .66 (.41) | .56 (.43) | 1.06 | .29 |
| Q learning rate ($\alpha_Q$) | .29 (.31) | .23 (.32) | .86 | .39 |
| Mixing parameter (m) | .64 (.34) | .48 (.39) | 2.04 | .04 |
| Inverse temperature ($\beta$) | .34 (.41) | .29 (.42) | .50 | .62 |

B = inverse temperature multiplied by 100

**Figure 4.**
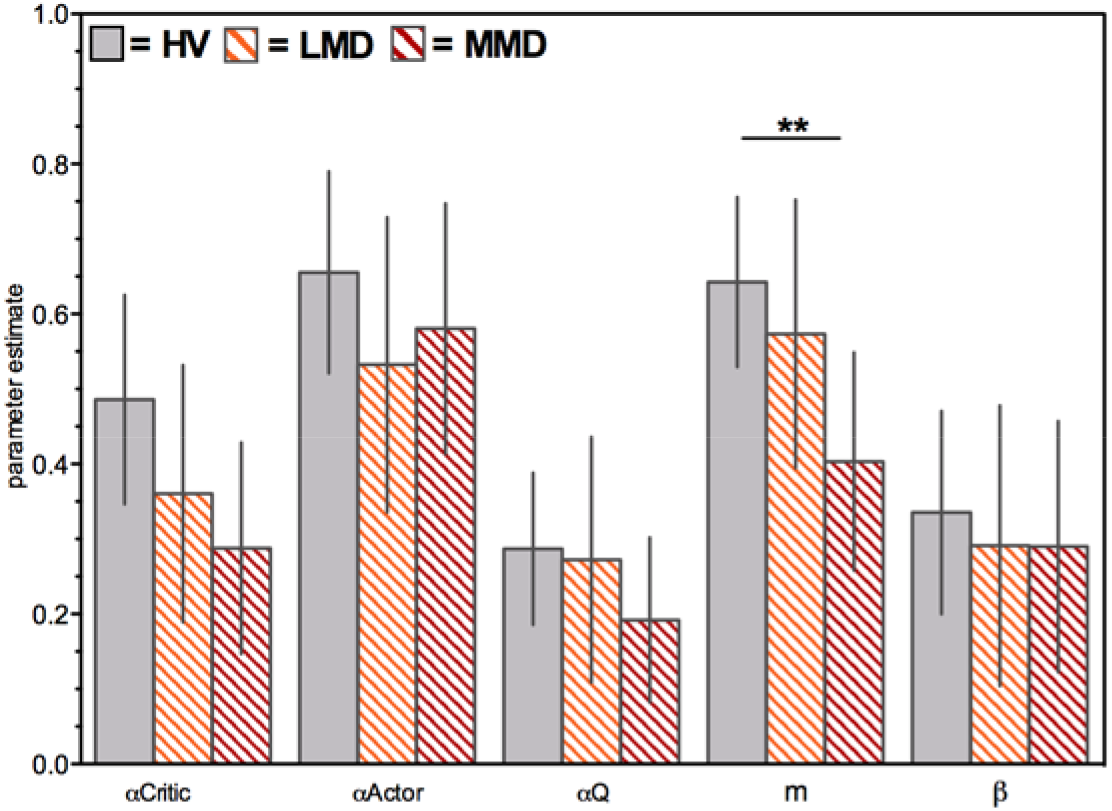
Hybrid-probability model parameters for HV, LMD and MMD. **p<.01

As predicted, the mixing parameter highly correlated with the value difference tracking slope in the entire sample, suggesting that greater reliance on Q-learning, in which choice values converge to their true expected values (16), was associated with better probability-magnitude integration (Spearman’s rho=.39, *p*<.001). The mixing parameter additionally correlated with the difference between performance on magnitude- and probability-discrimination trials (Spearman’s rho=.38, *p*<.001). This finding is line with our demonstration of the actor-critic being insensitive to reward magnitude (see “Model Selection and Comparison”) and suggests that better discrimination of reward magnitude was associated with more EV-based learning.

All in all, these results suggest that increased reliance on stimulus-response associations/decreased use of EV in PSZ with motivational deficits, as demonstrated by our hybrid-probability model, is associated with impaired probability-magnitude integration and may be driven by reduced sensitivity to reward magnitude specifically.

### Hybrid-probability model simulations

Hybrid-probability model Learning Phase simulations closely approximated empirical data, with a predicted main effect of group on accuracy for 90-10/3 (t_(85)_=3.21, *p*=.002), 80-20/2 (t_(85)_=3.52, *p*<.001), 80-20/1 (t_(85)_=2.28, *p*=.03) trials and a similar non-significant trend for 90-10/1 (t_(85)_=1.61, *p*=.11) trials (Figure 5A). Analyses of simulated data revealed either significant between-group differences, or strong trends toward significant between-group differences on all acquisition pair Test/Transfer trials (90-10/3 t_(85)_=2.32, *p*=.02; 80-20/2 t_(85)_=1.85, *p*=.07; 90-10/1 t_(85)_=2.41, *p*=.02; 80-20/1 t(85)=1.89, *p*=.06; Figure 5A; n simulations=20).

**Figure 5.**
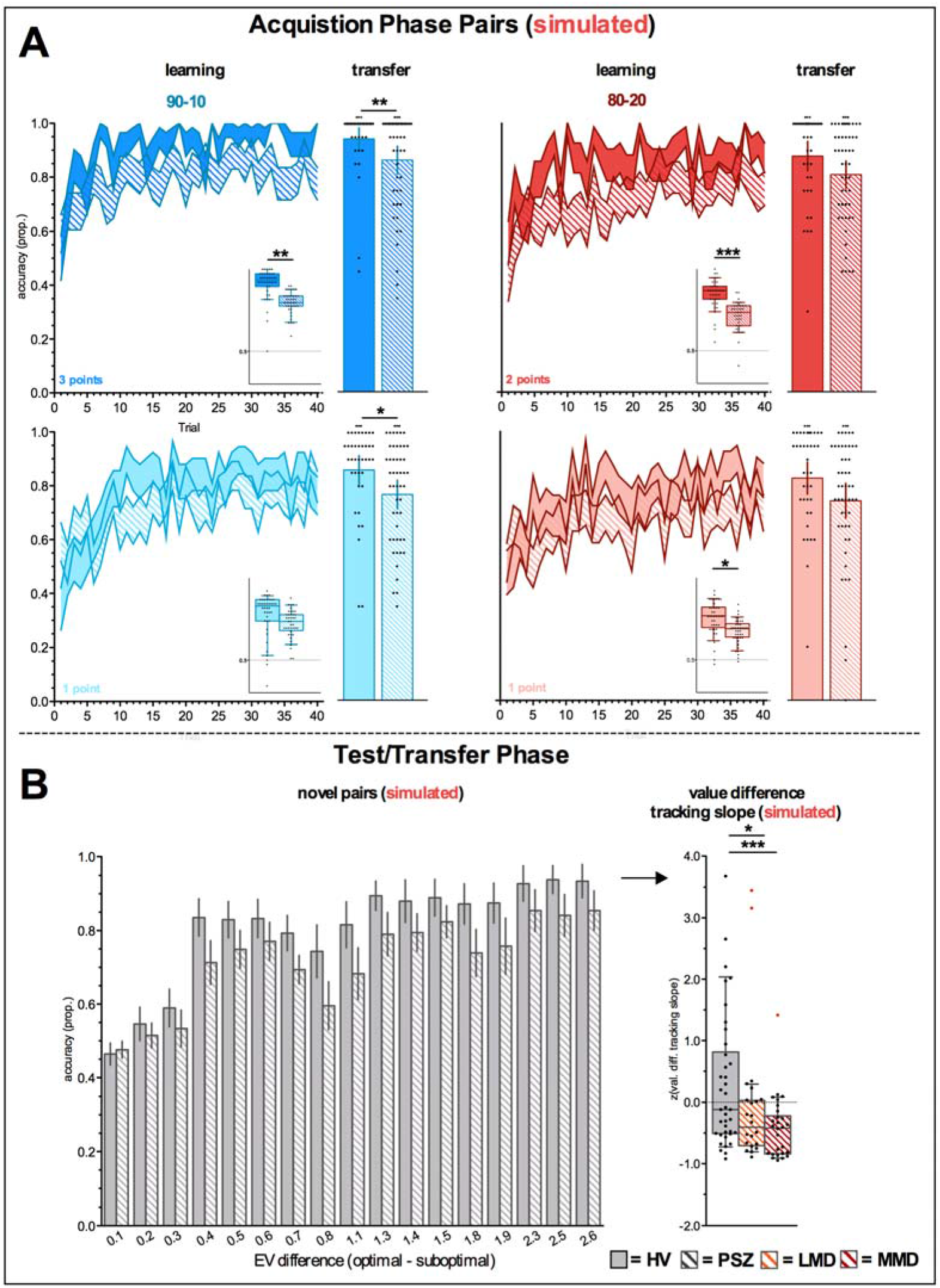
Hybrid-probability model simulations. **5A** Simulated Performance on Acquisition Phase Pairs during the Learning and Test/Transfer Phase. **5B** Simulated Test/Transfer phase performance on novel pairs presented for every value difference separately (“novel pairs”) and the value difference tracking (2 LMD and 1 MMD outliers marked in red). *p<.05, **p<.01, ***p<.001. Small asterisk above bars represents significance against chance.

Consistent with the empirical Test/Transfer Phase data, simulations revealed numerically greater deficits in PSZ for *easier* choices (greater EV difference). Greater deficits on trials on which probability-magnitude integration makes choices *easier* were also observed in PSZ relative to HV (e.g. 90/3-80/1 and 90/3-20/1 [“1.9” and “2.5” in Figure 5B]). Crucially, the hybrid-probability model predicted a numerically smaller value difference tracking slope in MMD (HV vs. LMD t_(56)_=.02, *p*=.2.44; HV vs. MMD t_(62)_=3.44, *p*=.001; LMD vs. HMD t_(44)_=1.29, *p*=.20; Figure 5B). Taken together, the hybrid-probability model could account for key aspects of the Learning and Test/Transfer Phase data, as well as highly specific performance deficits in PSZ with motivational deficits.

Associations with antipsychotic dosage and subgroup analyses are reported in the Supplemental Results.

## DISCUSSION

Our primary aim was to provide new mechanistic understanding of impaired probability-magnitude integration in PSZ with motivational deficits. Using computational models of RL, we demonstrate that a validated framework that emphasizes a reduction in EV-based decision-making and a, potentially compensatory, overutilization of stimulus-response associations can account for such impairments.

In the current experiment, successful integration of reward probability and magnitude decreased choice difficulty via the EV difference between two competing stimuli. PSZ were specifically impaired on trials with greater objective EV difference between two stimuli, as evidenced by the group difference in the Test/Transfer Phase value difference tracking slope, which was primarily driven by PSZ with motivational deficits. This observed inability to combine reward magnitude and probability in generating adaptive estimates of EV is in line with a large body of previous work, including findings of performance deficits in PSZ on the Iowa Gambling Task (10-12). Moreover, our results parallel those found in the effort literature, where motivational deficits are associated with abnormal effort cost computations (30), which are most pronounced in conditions with high reward magnitude/probability (31, 32). Here, we demonstrate that such deficits in PSZ extend to probabilistic RL paradigms. This is also a conceptual replication of a previous study in which we reported greater impairments for easier choices in PSZ during an RL paradigm in which reward probability determined stimulus EV (7). Altogether, this reconfirms the notion that performance deficits in PSZ increase with demands placed on putative prefrontal processes involved in EV estimation, which seems to be a rather robust finding in this population (6, 7, 27, 33) and may serve as one explanation for a reduced motivational drive.

Multiple mechanisms underlying EV estimation deficits in PSZ with motivational deficits have been proposed previously, including, but not limited to, insensitivity to gains (6, 34), a reduced ability to update expectations in response to positive RPEs (33, 35, 36), and susceptibility to context-dependent learning (7). Crucially, the hybrid computational framework that has been used to explain these deficits (6, 7) can additionally account for performance deficits in PSZ observed in the current experiment. Specifically, outcome probability-magnitude integration deficits in PSZ were characterized by increased reliance on value-less stimulus-associations (actor-critic), in lieu of EV-based decision-making (Q-learning). As predicted, individual Test/Transfer Phase value difference tracking slopes correlated significantly with estimates of individual mixing parameters - which capture the balance between Q- and actor-critic-type learning - suggesting a systematic relationship between EV-based learning and probability-magnitude integration. Importantly, individual mixing parameters correlated significantly (negatively) with motivational deficit severity, providing formal computational modeling evidence that impaired probability-magnitude integration in PSZ with motivational deficits arises from overutilization of stimulus-response associations. To our knowledge, this is the third report of decreased Q-/increased actor-critic-type learning in PSZ (with motivational deficits). The observation that various manifestations of disrupted processes in EV estimation coalesce in this single computational framework increases its generalizability and suggests potential for the computational hybrid model as a diagnostic tool for the detection of motivational deficits.

Outcome probability-magnitude integration deficits seemed to be driven by underutilization of reward magnitude specifically. This interpretation is supported by the between-group difference in performance improvement due to outcome magnitude versus probability increases (while holding the other constant). Moreover, this contrast (Figure 3) correlated with the value difference tracking slope in the entire sample, suggesting that better performance on magnitude trials was associated with more efficient probability-magnitude integration. This observation goes hand-in-hand with evidence for increased Q-learning in HV, a framework in which magnitude-driven RPEs also update the EV of a choice. Moreover, we demonstrated that magnitude was not used in the formation of response tendencies in the actor-critic architecture; the framework that could account for PSZ performance. Reduced reward (magnitude) sensitivity is in line with reports on anhedonia (37, 38), and observations in PSZ with motivational deficits during a outcome probability-magnitude integration task (14).

Speculating on the neural mechanisms involved, the OFC has consistently been associated with tracking of reward history and rapid EV updating (20, 39-41). Pre-clinical work suggests that reward magnitude is encoded by neurons in the basolateral amygdala and OFC (39, 42-45). Moreover, lesions to the basolateral amygdala alter reward magnitude encoding (46) and value-based decision-making (47-49) involving the OFC. In contrast, reward probability may be associated with increased midbrain (28, 29) and ventral striatum (45, 50, 51) activity, although the latter has also been reported to signal reward magnitude (45, 51). The specific inability to utilize reward magnitude information in combination with previously-reported EV deficits in PSZ may therefore point to a deficit in the basolateral amygdala-OFC projection or ventral striatum. Indeed, abnormal OFC and striatal reward signals have been reported in the psychosis spectrum (52-55), which occasionally travel with motivational deficit severity. Future studies aimed at disambiguating the neural signals associated with reward magnitude, reward probability, and probability-magnitude integration, may provide valuable insights into the nature of reward processing deficits in PSZ (with motivational deficits).

In conclusion, we provide formal evidence for the notion that failure to integrate reward probability and magnitude in PSZ with motivational deficits is associated with overreliance on the learning of value-less stimulus-response associations. Performance improvements associated with increases in reward magnitude, in combination with analyses using our computational hybrid model, suggests that such deficits may be specifically associated with impairment in the ability to precisely and adaptively represent reward magnitude. The results presented in this manuscript add to the generalizability of the computational hybrid model in capturing a broad range of EV estimation deficits in PSZ with motivational deficits.

## LIMITATIONS

Only a small number of trials were available to directly investigate probability and magnitude processing. Moreover, magnitude and probability were not fully orthogonalized in this experiment, slightly reducing the total number of trials that could be used to study reward probability. Importantly, however, the key aim of this study was to study probability-magnitude integration deficits in an RL context, rather than specific deficits in reward magnitude or probability processing. Moreover, a selective deficit in learning from reward magnitude is in line with previous work (14, 38), and was predicted and demonstrated by our computational modeling framework. Finally, our sample consisted of chronic, medicated PSZ. While antipsychotic drug doses were not associated with any outcome measures, a follow-up investigation in antipsychotic-naïve PSZ could provide more information on symptom-specificity.

## REFERENCES

1. Strauss GP, Waltz JA, Gold JM (2014): A review of reward processing and motivational impairment in schizophrenia. Schizophrenia bulletin. 40 Suppl 2:S107–116.

2. Fervaha G, Foussias G, Agid O, Remington G (2015): Motivational deficits in early schizophrenia: prevalent, persistent, and key determinants of functional outcome. Schizophrenia research. 166:9–16.

3. Dickinson D, Bellack AS, Gold JM (2007): Social/communication skills, cognition, and vocational functioning in schizophrenia. Schizophrenia bulletin. 33:1213–1220.

4. Velligan DI, Kern RS, Gold JM (2006): Cognitive rehabilitation for schizophrenia and the putative role of motivation and expectancies. Schizophrenia bulletin. 32:474–485.

5. Fervaha G, Foussias G, Agid O, Remington G (2014): Motivational and neurocognitive deficits are central to the prediction of longitudinal functional outcome in schizophrenia. Acta Psychiatr Scand. 130:290–299.

6. Gold JM, Waltz JA, Matveeva TM, Kasanova Z, Strauss GP, Herbener ES, et al. (2012): Negative symptoms and the failure to represent the expected reward value of actions: behavioral and computational modeling evidence. Archives of general psychiatry. 69:129–138.

7. Hernaus D, Gold JM, Waltz JA, Frank MJ (2018): Impaired Expected Value Computations Coupled With Overreliance on Stimulus-Response Learning in Schizophrenia. Biological psychiatry Cognitive neuroscience and neuroimaging.

8. Waltz JA, Gold JM (2016): Motivational Deficits in Schizophrenia and the Representation of Expected Value. Current topics in behavioral neurosciences. 27:375–410.

9. Bechara A, Damasio AR, Damasio H, Anderson SW (1994): Insensitivity to future consequences following damage to human prefrontal cortex. Cognition. 50:7–15.

10. Brown EC, Hack SM, Gold JM, Carpenter WT, Jr., Fischer BA, Prentice KP, et al. (2015): Integrating frequency and magnitude information in decision-making in schizophrenia: An account of patient performance on the Iowa Gambling Task. Journal of psychiatric research. 66-67:16–23.

11. Brambilla P, Perlini C, Bellani M, Tomelleri L, Ferro A, Cerruti S, et al. (2013): Increased salience of gains versus decreased associative learning differentiate bipolar disorder from schizophrenia during incentive decision making. Psychological medicine. 43:571–580.

12. Kim MS, Kang BN, Lim JY (2016): Decision-making deficits in patients with chronic schizophrenia: Iowa Gambling Task and Prospect Valence Learning model. Neuropsychiatric disease and treatment. 12:1019–1027.

13. Brown JK, Waltz JA, Strauss GP, McMahon RP, Frank MJ, Gold JM (2013): Hypothetical decision making in schizophrenia: the role of expected value computation and “irrational” biases. Psychiatry research. 209:142–149.

14. Albrecht MA, Waltz JA, Frank MJ, Gold JM (2016): Probability and magnitude evaluation in schizophrenia. Schizophrenia research Cognition. 5:41–46.

15. Kasanova Z, Waltz JA, Strauss GP, Frank MJ, Gold JM (2011): Optimizing vs. matching: response strategy in a probabilistic learning task is associated with negative symptoms of schizophrenia. Schizophrenia research. 127:215–222.

16. Watkins C, Dayan P (1992): Q-learning. Mach Learning.279–292.

17. Furuyashiki T, Gallagher M (2007): Neural encoding in the orbitofrontal cortex related to goal-directed behavior. Annals of the New York Academy of Sciences. 1121:193–215.

18. Padoa-Schioppa C, Cai X (2011): The orbitofrontal cortex and the computation of subjective value: consolidated concepts and new perspectives. Annals of the New York Academy of Sciences. 1239:130–137.

19. Rich EL, Wallis JD (2016): Decoding subjective decisions from orbitofrontal cortex. Nature neuroscience. 19:973–980.

20. Padoa-Schioppa C, Assad JA (2006): Neurons in the orbitofrontal cortex encode economic value. Nature. 441:223–226.

21. Gottfried JA, O’Doherty J, Dolan RJ (2003): Encoding predictive reward value in human amygdala and orbitofrontal cortex. Science. 301:1104–1107.

22. Rolls ET, McCabe C, Redoute J (2008): Expected value, reward outcome, and temporal difference error representations in a probabilistic decision task. Cerebral cortex. 18:652–663.

23. Rescorla RA, Wagner AR (1972): in Classical Conditioning II: Current Research and Theory, eds Black AH, Prokasy WF. New York City, NY: Appleton–Century Crofts.

24. Joel D, Niv Y, Ruppin E (2002): Actor-critic models of the basal ganglia: new anatomical and computational perspectives. Neural networks: the official journal of the International Neural Network Society. 15:535–547.

25. Collins AG, Frank MJ (2014): Opponent actor learning (OpAL): modeling interactive effects of striatal dopamine on reinforcement learning and choice incentive. Psychological review. 121:337–366.

26. Frank MJ, Claus ED (2006): Anatomy of a decision: striato-orbitofrontal interactions in reinforcement learning, decision making, and reversal. Psychological review. 113:300–326.

27. Waltz JA, Frank MJ, Robinson BM, Gold JM (2007): Selective reinforcement learning deficits in schizophrenia support predictions from computational models of striatal-cortical dysfunction. Biological psychiatry. 62:756–764.

28. Bayer HM, Glimcher PW (2005): Midbrain dopamine neurons encode a quantitative reward prediction error signal. Neuron. 47:129–141.

29. Bayer HM (2004): A role for the substantia nigra in learning and motor control. New York: New York University.

30. Culbreth AJ, Moran EK, Barch DM (2018): Effort-Based Decision-Making in Schizophrenia. Current opinion in behavioral sciences. 22:1–6.

31. Treadway MT, Peterman JS, Zald DH, Park S (2015): Impaired effort allocation in patients with schizophrenia. Schizophrenia research. 161:382–385.

32. Gold JM, Strauss GP, Waltz JA, Robinson BM, Brown JK, Frank MJ (2013): Negative symptoms of schizophrenia are associated with abnormal effort-cost computations. Biological psychiatry. 74:130–136.

33. Dowd EC, Frank MJ, Collins A, Gold JM, Barch DM (2016): Probabilistic Reinforcement Learning in Patients With Schizophrenia: Relationships to Anhedonia and Avolition. Biological psychiatry Cognitive neuroscience and neuroimaging. 1:460–473.

34. Hartmann-Riemer MN, Aschenbrenner S, Bossert M, Westermann C, Seifritz E, Tobler PN, et al. (2017): Deficits in reinforcement learning but no link to apathy in patients with schizophrenia (vol 7, 40352, 2017). Scientific reports. 7.

35. Waltz JA, Xu Z, Brown EC, Ruiz RR, Frank MJ, Gold J (2017): Motivational Deficits in Schizophrenia Are Associated With Reduced Differentiation Between Gain and Loss-Avoidance Feedback in the Striatum. Biological Psychiatry: CNNI.

36. Deserno L, Heinz A, Schlagenhauf F (2017): Computational approaches to schizophrenia: A perspective on negative symptoms. Schizophrenia research. 186:46–54.

37. Huys QJ, Pizzagalli DA, Bogdan R, Dayan P (2013): Mapping anhedonia onto reinforcement learning: a behavioural meta-analysis. Biology of mood & anxiety disorders. 3:12.

38. Vignapiano A, Mucci A, Ford J, Montefusco V, Plescia GM, Bucci P, et al. (2016): Reward anticipation and trait anhedonia: An electrophysiological investigation in subjects with schizophrenia. Clin Neurophysiol. 127:2149–2160.

39. Burke SN, Thome A, Plange K, Engle JR, Trouard TP, Gothard KM, et al. (2014): Orbitofrontal cortex volume in area 11/13 predicts reward devaluation, but not reversal learning performance, in young and aged monkeys. J Neurosci. 34:9905–9916.

40. Riceberg JS, Shapiro ML (2017): Orbitofrontal Cortex Signals Expected Outcomes with Predictive Codes When Stable Contingencies Promote the Integration of Reward History. J Neurosci. 37:2010–2021.

41. Murray EA, Moylan EJ, Saleem KS, Basile BM, Turchi J (2015): Specialized areas for value updating and goal selection in the primate orbitofrontal cortex. eLife. 4.

42. Smith BW, Mitchell DG, Hardin MG, Jazbec S, Fridberg D, Blair RJ, et al. (2009): Neural substrates of reward magnitude, probability, and risk during a wheel of fortune decision-making task. NeuroImage. 44:600–609.

43. Bermudez MA, Schultz W (2010): Reward magnitude coding in primate amygdala neurons. Journal of neurophysiology. 104:3424–3432.

44. Saez RA, Saez A, Paton JJ, Lau B, Salzman CD (2017): Distinct Roles for the Amygdala and Orbitofrontal Cortex in Representing the Relative Amount of Expected Reward. Neuron. 95:70–77 e73.

45. Stoppel CM, Boehler CN, Strumpf H, Heinze HJ, Hopf JM, Schoenfeld MA (2011): Neural processing of reward magnitude under varying attentional demands. Brain research. 1383:218–229.

46. Rudebeck PH, Mitz AR, Chacko RV, Murray EA (2013): Effects of amygdala lesions on reward-value coding in orbital and medial prefrontal cortex. Neuron. 80:1519–1531.

47. Lichtenberg NT, Pennington ZT, Holley SM, Greenfield VY, Cepeda C, Levine MS, et al. (2017): Basolateral Amygdala to Orbitofrontal Cortex Projections Enable Cue-Triggered Reward Expectations. J Neurosci. 37:8374–8384.

48. Fiuzat EC, Rhodes SE, Murray EA (2017): The Role of Orbitofrontal-Amygdala Interactions in Updating Action-Outcome Valuations in Macaques. J Neurosci. 37:2463–2470.

49. Rudebeck PH, Ripple JA, Mitz AR, Averbeck BB, Murray EA (2017): Amygdala Contributions to Stimulus-Reward Encoding in the Macaque Medial and Orbital Frontal Cortex during Learning. J Neurosci. 37:2186–2202.

50. Abler B, Walter H, Erk S, Kammerer H, Spitzer M (2006): Prediction error as a linear function of reward probability is coded in human nucleus accumbens. NeuroImage. 31:790–795.

51. Yacubian J, Sommer T, Schroeder K, Glascher J, Braus DF, Buchel C (2007): Subregions of the ventral striatum show preferential coding of reward magnitude and probability. NeuroImage. 38:557–563.

52. Segarra N, Metastasio A, Ziauddeen H, Spencer J, Reinders NR, Dudas RB, et al. (2016): Abnormal Frontostriatal Activity During Unexpected Reward Receipt in Depression and Schizophrenia: Relationship to Anhedonia. Neuropsychopharmacology: official publication of the American College of Neuropsychopharmacology. 41:2001–2010.

53. de Leeuw M, Kahn RS, Vink M (2015): Fronto-striatal dysfunction during reward processing in unaffected siblings of schizophrenia patients. Schizophrenia bulletin. 41:94–103.

54. Waltz JA, Schweitzer JB, Gold JM, Kurup PK, Ross TJ, Salmeron BJ, et al. (2009): Patients with schizophrenia have a reduced neural response to both unpredictable and predictable primary reinforcers. Neuropsychopharmacology: official publication of the American College of Neuropsychopharmacology. 34:1567–1577.

55. Radua J, Schmidt A, Borgwardt S, Heinz A, Schlagenhauf F, McGuire P, et al. (2015): Ventral Striatal Activation During Reward Processing in Psychosis: A Neurofunctional Meta-Analysis. JAMA psychiatry. 72:1243–1251.

